# *Pseudozyma* saprotrophic yeasts have retained a large effector arsenal, including functional Pep1 orthologs

**DOI:** 10.1101/489690

**Authors:** Rahul Sharma, Bilal Ökmen, Gunther Doehlemann, Marco Thines

**Affiliations:** Senckenberg BiK-F, Frankfurt am Main, Germany; Goethe University, Biosciences, Institute of Ecology, Evolution and Diversity, Frankfurt am Main, Germany; Botanical Institute and CEPLAS, University of Cologne, BioCenter, Cologne, Germany; Integrative Fungal Research Cluster (IPF), Frankfurt am Main, Germany

**Keywords:** core effectors, effector complementation, plant pathogens, *Pseudozyma*, *Ustilago*, yeast

## Abstract

The basidiomycete smut fungi are predominantly plant parasitic, causing severe losses in some crops. Most species feature a saprotrophic haploid yeast stage, and several smut fungi are only known from this stage, with some isolated from habitats without suitable hosts, e.g. from Antarctica. Thus, these species are generally believed to be apathogenic, but recent findings that some of these might have a plant pathogenic sexual counterpart, casts doubts on the validity of this hypothesis. Here, four *Pseudozyma* genomes were re-annotated and compared to published smut pathogens and the well-characterised effector gene *Pep1* from these species was checked for its ability to complement a *Pep1* deletion strain of *Ustilago maydis*. It was found that 113 high-confidence putative effector proteins were conserved among smut and *Pseudozyma* genomes. Among these were several validated effector proteins, including Pep1. By genetic complementation we show that *Pep1* homologs from the supposedly apathogenic yeasts restore virulence in *Pep1*-deficient mutants *Ustilago maydis*. Thus, it is concluded that *Pseudozyma* species have retained a suite of effectors. This hints at the possibility that *Pseudozyma* species have kept an unknown plant pathogenic stage for sexual recombination or that these effectors have positive effects when colonising plant surfaces.

## Introduction

Smut fungi in a broad sense are one of the three major lineages of Basidiomycota. Some species threaten crop plants, e.g. *Ustilago maydis* (maize) and *Ustilago scitaminea* (sugarcane). Smut fungi are dimorphic, with a filamentous and a yeast-like morph in different life stages. The yeast stage is haploid and saprotrophic, the hyphal stage is dicaryotic and obligate plant parasitic. Yeasts of matching sex form fusing conjugation hyphae, producing an infective dikaryon penetrating the host or colonisation. However, some smuts are known only from their yeast stage. Since the ending of dual naming for anamorphic and teleomorphic fungi (Hawksworth *et al.*, 2011), the yeast genus *Pseudozyma* became obsolete, being scattered throughout the smut phylogeny (Wang *et al.*, 2015). It was recently established that *Pseudozyma prolifica* is conspecific with *U. maydis* (Wang *et al.*, 2015), but it is believed that other *Pseudozyma* species have lost pathogenicity and exist only as yeasts (Lefebvre *et al.*, 2013; Morita *et al.*, 2013; Morita *et al.*, 2014). A well-known species of these is the biocontrol agent *Pseudozyma aphidis* (Gafni *et al.*, 2015), closely related or probably conspecific with *Moesziomyces bullatus* (Kruse *et al.*, 2017).

A key to investigating the apathogenic or pathogenic nature of *Pseudozyma*-like yeasts are putative secreted effector proteins (PSEPs). For successful colonization of hosts, pathogens secrete hundreds of effector proteins (Latijnhouwers *et al.*, 2003). Effectors often have limited sequence conservation (Schirawski *et al.*, 2010; Kemen *et al.*, 2011; Sharma *et al.*, 2014), but some are conserved in genomes of related pathogens (Sharma *et al.* 2014, 2015; Quinn *et al.* 2013; Hemetsberger *et al.* 2015; Lanver et al. (2017). Previous definitions of putative effector proteins were often associated with several restrictions, such as size cutoff (e.g. below 300aa), certain amino acid composition (e.g. Cys-rich), or the absence of sequence motifs predicting an enzymatic function. However, such formal restrictions artificially exclude a large fraction of proteins that contribute to virulence and interfere with the plant immune system. Effectors with enzymatic functions have actually been identified in various filamentous plant pathogens (Franceschetti *et al.*, 2017). Recently, the *U. maydis* metalloprotease Fly1 was found to contribute to virulence by targeting maize chitinase. Here, we investigated conservation of PSEPs, including effectors with predicted functional domains, among *Pseudozyma* genomes (Morita *et al.*, 2013; Konishi *et al.*, 2013; Oliveira *et al.*, 2013; Lorenz *et al.*, 2014) to evaluate the potential pathogenicity of *Pseudozyma*-like yeasts. We were focusing on well-studied effectors, in particular the core-effector *Pep1*, which previously was identified as an essential virulence factor of *U. maydis* which is functionally conserved amongst pathogenic smuts (Hemetsberger *et al.*, 2012, 2015).

## Results

Of 211 core PSEPs, a total of 178, 199, 182 and 171 candidates were conserved in the non-pathogenic yeasts *P. antarctica, P. hubeiensis, P. brasiliensis*, and *P. aphidis*, respectively (Tables 1, 2). Of these, 158, 182, 166, and 151, respectively, were predicted to have a secretion signal (Table 2). In total, 113 PSEPs were found to be conserved among all eight genomes. Functional annotation revealed features associated with pathogenicity, such as glycoside hydrolase and aspartic peptidase domains (Supplementary Table 2).

**Table 1.** Conservation of functionally characterized pathogenicity related proteins of *U. maydis* among four *Pseudozyma* genomes.

| *U. maydis* gene | Gene name | *P. antarctica* | *P. aphidis* | *P. brasiliensis* | *P. hubeiensis* |
| --- | --- | --- | --- | --- | --- |
| UMAG_04433 | *Hum3* | Pant_3371_t | Paph_3632_t.1* | Pbra_985_t | Phub_3033_t |
| UMAG_01987 | *Pep1* | Pant_5912_t | Paph_3137_t | Pbra_3010_t | Phub_108_t |
| UMAG_01375 | *Pit2* | Pant_3484_t*** | Paph_3237_t*** | Pbra_3853_t*** | Phub_3768_t |
| UMAG_05302 | *Tin2* | Pant_4343_t.2*** | Paph_1304_t.2*** | Pbra_3373_t | Phub_4666_t.1** |
| UMAG_05731 | *Cmu1* | Pant_6209_t | Paph_4714_t | Pbra_2221_t | Phub_1879_t |

\* Sequence contains a stretch of Ns, could be resolved by experimental methods. \*\* No signal peptide predicted by using SignalP4.1, but by TargetP1 and Protcomp9. Genes with secretion signal and some conservation of some domains, but not matching the strict orthology criteria over the entire length.

**Table 2.** Number of orthologs of putative effectors found among the four *Pseudozyma* genomes.

| Species | All Orthologs | Orthologs with SP in all species |
| --- | --- | --- |
| *P. antarctica* | 178 | 158 |
| *P. hubeiensis* | 199 | 182 |
| *P. brasiliensis* | 182 | 166 |
| *P. aphidis* | 171 | 151 |

Almost all of the well-studied *Ustilago maydis* virulence factors conserved among Ustilaginales were found conserved in the four *Pseudozyma* genomes (Table 1). The conservation of the *Pit* cluster (Doehlemann *et al.*, 2011) *Cmu1* (Djamei *et al.*, 2011), *Cwh41* (Martínez-Soto *et al.*, 2013), and *Hum3* (Muller *et al.*, 2008) was found in all four *Pseudozyma* genomes (Table 1). *Tin2* (Tanaka *et al.*, 2014) was also found conserved, but the *P. hubeiensis* ortholog was lacking a strong secretion signal (Table 1). The *Pep1* effector was found to be conserved among the eight genomes, with all structural features intact (Figure 1). Besides these effectors with already known virulence function, also the membrane proteins Msb2 (Lanver *et al.*, 2010) and Pit1 (Doehlemann et al., 2011), which hold crucial virulence functions, but are also known from non-pathogenic species were conserved amongst the yeast genomes.

**Figure 1.**
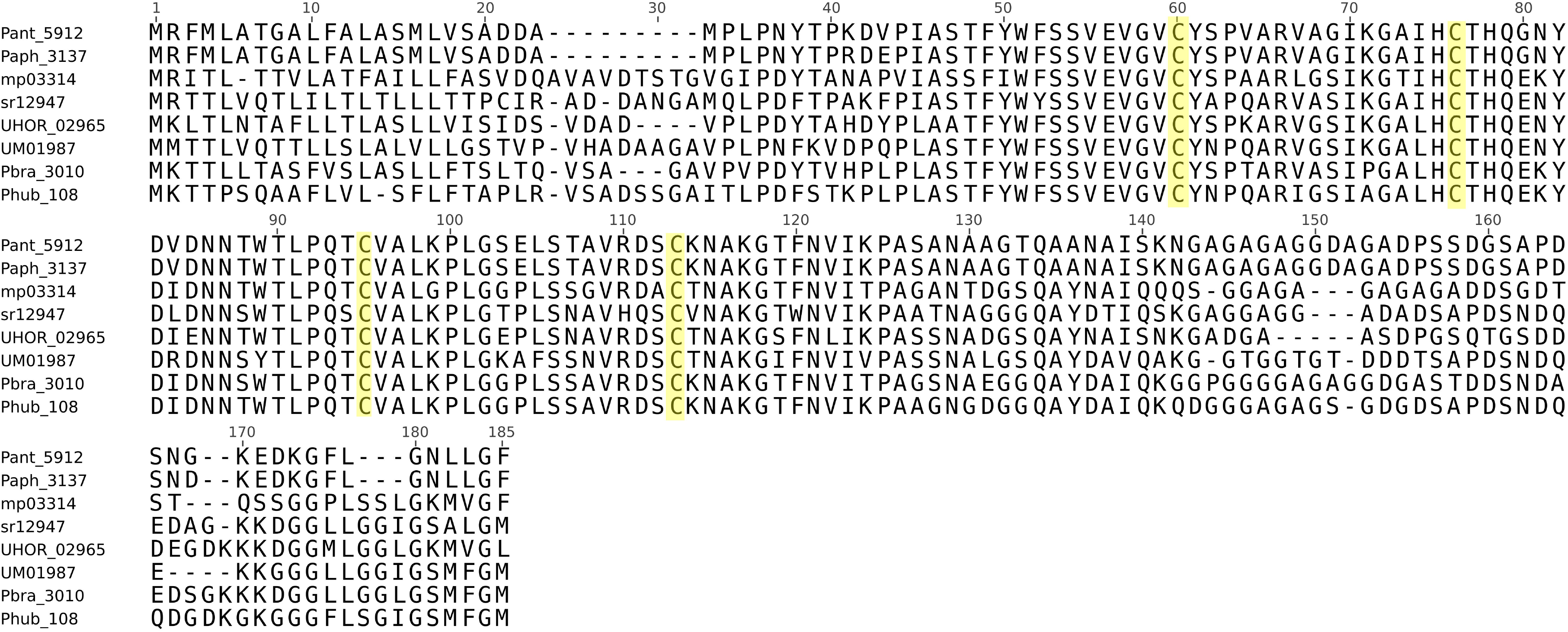
Multiple sequence alignment of eight candidate *Pep1* proteins from pathogens and *Pseudozyma* yeasts. Multiple sequence alignment (MSA) shows high sequence conservation of the candidate *Pep1* effector proteins. The conserved four cysteine residues needed for the funtion of *Pep1* are highlighted in yellow.

To test the functionality as virulence factors on the example of the well-studied effector Pep1, coding regions of the *Pseudozyma Pep1* orthologs were fused to the *U. maydis pep1* promoter and stably integrated in the *ip*-locus of *U. maydis* strain SG200Δpep1, which is unable to infect maize plants due to the deletion of *Pep1* (Doehlemann *et al.*, 2009). The resulting *U. maydis* strains were verified by Southern Blot (Supplementary Figure 1) and subsequently inoculated to maize seedlings. Strikingly, scoring of *U. maydis* tumour formation revealed that all strains expressing Pep1 homologs from *Pseudozyma* species were fully virulent, i.e. produced plant tumours were indistinguishable from the progenitor strain SG200 (Figure 2). This result demonstrates that *Pseudozyma* species encode functional orthologs of the Pep1 effector, which restore virulence of *U. maydis* in maize.

**Figure 2.**
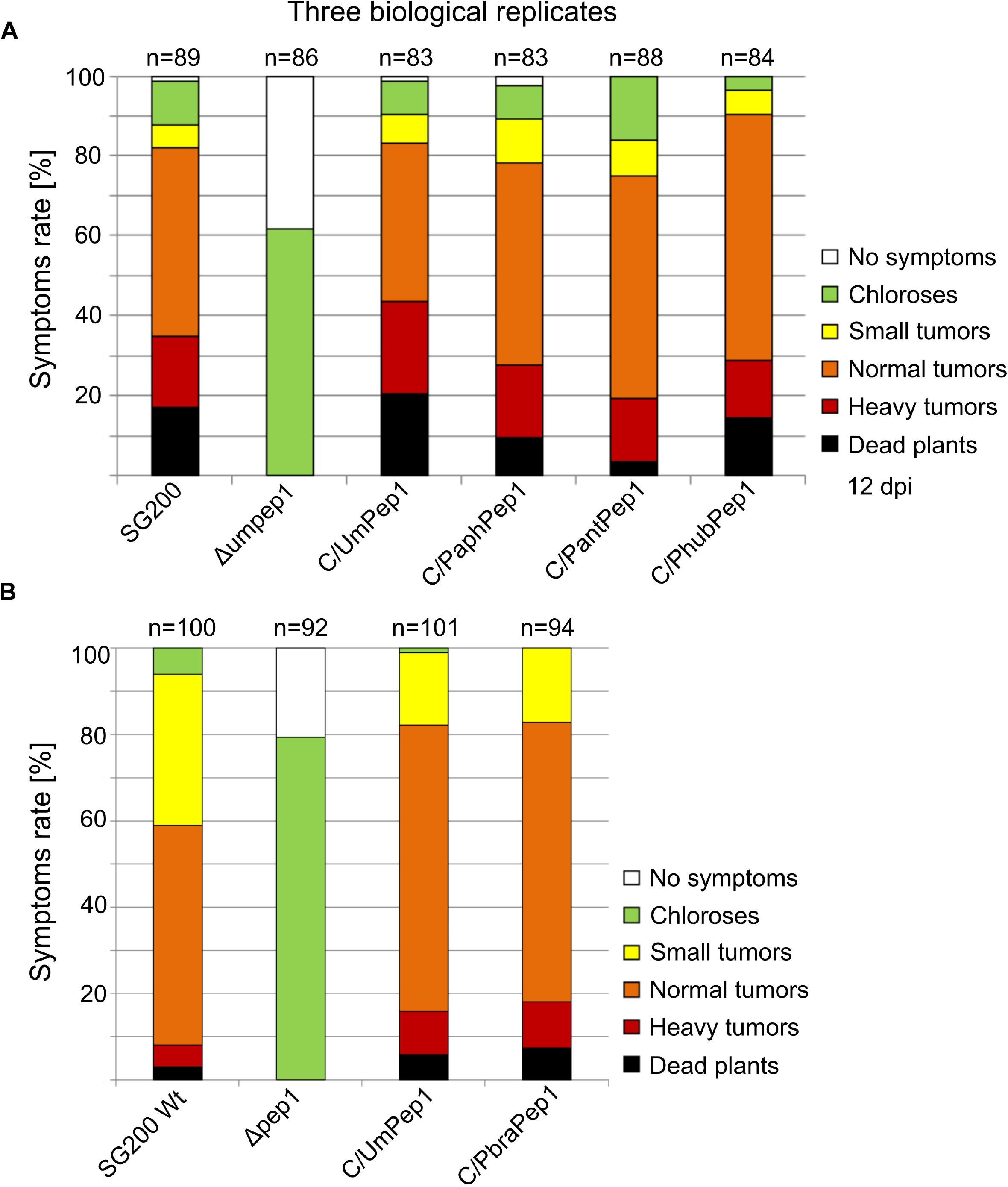
Disease assay on EGB maize lines. For testing biological conservation, Pep1 orthologs from *Pseudozyma* yeasts were expressed in a *Pep1* deletion background (SG200Δumpep1). The restoration of pathogenicity in the complemented lines is indicative of functional conservation.

## Discussion

To avoid recognition by R-proteins, many secreted effectors adapt quickly and show limited conservation (Schirawski *et al.*, 2010; Laurie *et al.*, 2012). However, conservation of putative secreted effector protein (PSEP)-encoding genes among related pathogens has been reported (Hemetsberger *et al.*, 2015; Sharma et al., 2015). These effectors have been termed “core” effectors (Sharma et al. 2014). The well-studied Pep1 effector, required for successful host colonisation (Doehlemann *et al.*, 2009), is an archetypal core effector, functionally conserved among monocot and dicot infecting smut pathogens (Hemetsberger *et al.*, 2015). Our finding that also Cmu1, Tin2, and Hum3 remained conserved with secretion signal peptides in the genomes of smuts fungi only known from the yeast stage suggests that they either feature an unknown pathogenic stage, or that effectors also have a positive effect when settling on plant surfaces. Also in *P. antarctica* isolated from sediments in Antarctica, these effectors remained conserved, with secretion signal peptides intact. If these effectors were not needed anymore, it can be expected that they would either get lost quickly, as in *U. pennsylvanica* after a host jump from monocots to dicots (Sharma *et al.*, 2014), or acquire new functions (Sharma *et al.*, 2015). For *Pep1*, the finding that orthologs from all yeast-only species were able to fully restore pathogenicity in *U. maydis*, demonstrates that its virulence function remained conserved amongst species. Moreover, our finding highlights that more research is needed to investigate, if fully saprotrophic species of the Ustilaginales exist (Kruse *et al.*, 2017) and if so, how different lifestyles of smut yeasts evolved. Regarding the functional conservation of Pep1 one could speculate that its function to suppress PAMP-triggered ROS generation also benefits epiphytic yeasts on plant surfaces. However, in *U. maydis* transcription of Pep1, as well as that of other known effectors, is only activated upon mating when compatible heterodimers of the b-transcription factor are present in the cell. Therefore, a putative role of effectors in non-biotrophic stages would imply a fundamental change in the transcriptional regulatory cascade of anamorphic smut yeasts. Even though the alternative explanation that all the conserved proteins would also benefit a saprotrophic yeasts seems highly unlikely, as several are known to interact only with targets inside the plant cytoplasm. Given the conservation of more than 100 PSEPs and almost all core effectors with known virulence activity, it seems more likely that *Pseudozyma* species have a plant pathogenic stage, rather than that they have lost it only recently, simultaneously in four species. It is conceivable that those few species frequently encountered as yeasts, are competitive saprotrophs and that the plant pathogenic stage is only maintained to allow for infrequent sexual recombination.

## Materials and Methods

### Bioinformatics

Out of 248 PSEPs conserved among the four smut genomes i.e. *Ustilago maydis* (Kämper *et al.*, 2006), *U. hordei* (Laurie *et al.*, 2012), *U. reiliana* (Schirawski *et al.*, 2010), and *U. pennsylvanica* (Sharma *et al.*, 2014), a high-confidence (secretion strongly supported) core set of 211 PSEPs was inferred (Sharma *et al.*, 2015). The genomes of four *Pseudozyma* species, *P. antarctica, P. hubeiensis, P. brasiliensis*, and *P. aphidis*, were scanned for the presence of the 211 PSEPs, with *U. maydis* proteins as query. To investigate the conservation, also *ab initio* prediction was done using GeneMark (Ter-Hovhannisyan *et al.*, 2008), trained on the other Ustilaginales. The resulting protein sequences were aligned to the 211 PSEPs of *U. maydis* using Blastp. A PSEP was considered present if exceeding 45% identity, an e-value of e-5, and alignment coverage of 60%. Candidate orthologs were scanned for secretion signals as described before (Sharma *et al.*, 2015). Start-codon positions of candidate orthologs were manually checked and corrected, if necessary, using the well-annotated *U. maydis* proteins as reference.

The PSEPs conserved among smuts and yeasts were annotated based on *U. maydis* proteins (ftp://ftpmips.gsf.de/fungi/Ustilaginaceae/Ustilago_maydis_521/) and InterProScan (Quevillon *et al.*, 2005) using Blast2GO (Conesa *et al.*, 2005). Particular attention was paid to validated effector proteins of *U. maydis*.

### Δ*umpep1* complementation and disease assay

In order to show functional conservation between the *Pep1* orthologs, those from *P. antarctica, P. hubeiensis, P. brasiliensis*, and *P. aphidis* were amplified by PCR with the primers given in Supplementary Table 1 and subsequently expressed in the *U. maydis* SG200Δpep1 strain, which is deleted for *pep1* (Doehlemann *et al.*, 2009). For proper expression, *pep1* orthologs were expressed controlled by the endogenous *U. maydis pep1*-promoter, integrated in single copy (checked by Southern blots) into the *ip*-locus of *U. maydis*, as described previously (Hemetsberger *et al.*, 2015).

The *U. maydis* disease assays were performed according to Hemetsberger *et al.* (2015). Briefly, the cell culture control and complementation strains were inoculated onto maize seedlings (variety Early Golden Bantam) with a syringe and needle into the leaf whirl. All assays were performed in biological triplicates (≥30 plants each). The quantification of disease symptoms was performed at 12 dpi as described previously (Kämper *et al.*, 2006).

## Supporting information

## Acknowledgements

RS carried out bioinformatics work and participated in data analysis; BO carried out the molecular lab work, participated in data analysis, and edited the manuscript; GD designed the study, participated in data analysis and drafted the manuscript; MT conceived of the study, designed the study, participated in data analysis, coordinated the study and drafted the manuscript. This study was funded by the LOEWE initiative of the government of Hessen, in the framework of the cluster for Integrative Fungal Research (IPF) and the Center of Excellence on Plant Science (CEPLAS). We thank Melanie Kastl and Raphael Wemhöner for assistance in generation of *U. maydis* strains and plant infection assays.

## Supporting information figure caption

**Figure S1. Southern blot analysis to confirm single integration events.** All complementation events were performed in the *ip* locus in SG200Δumpep1 background. Restriction enzyme, DNA probe that were used and the expected fragments sizes for each southern blot analysis are indicated below each picture. Red arrows indicate single integration event in the correct genomic locus.

**Table S1.**
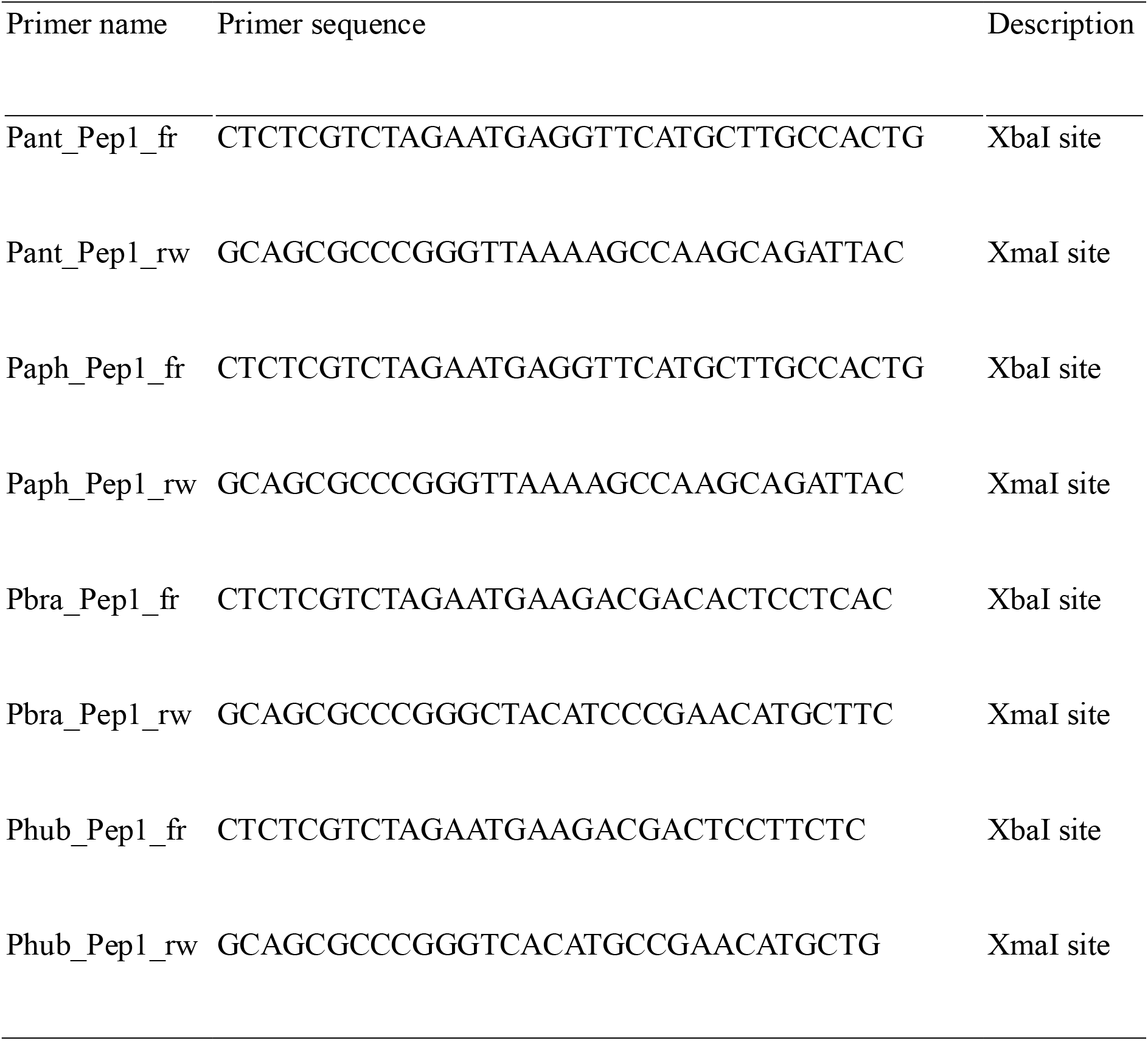
Primers for delta-umpep1 complementation with *Pep1* homologs

**Table S2.**
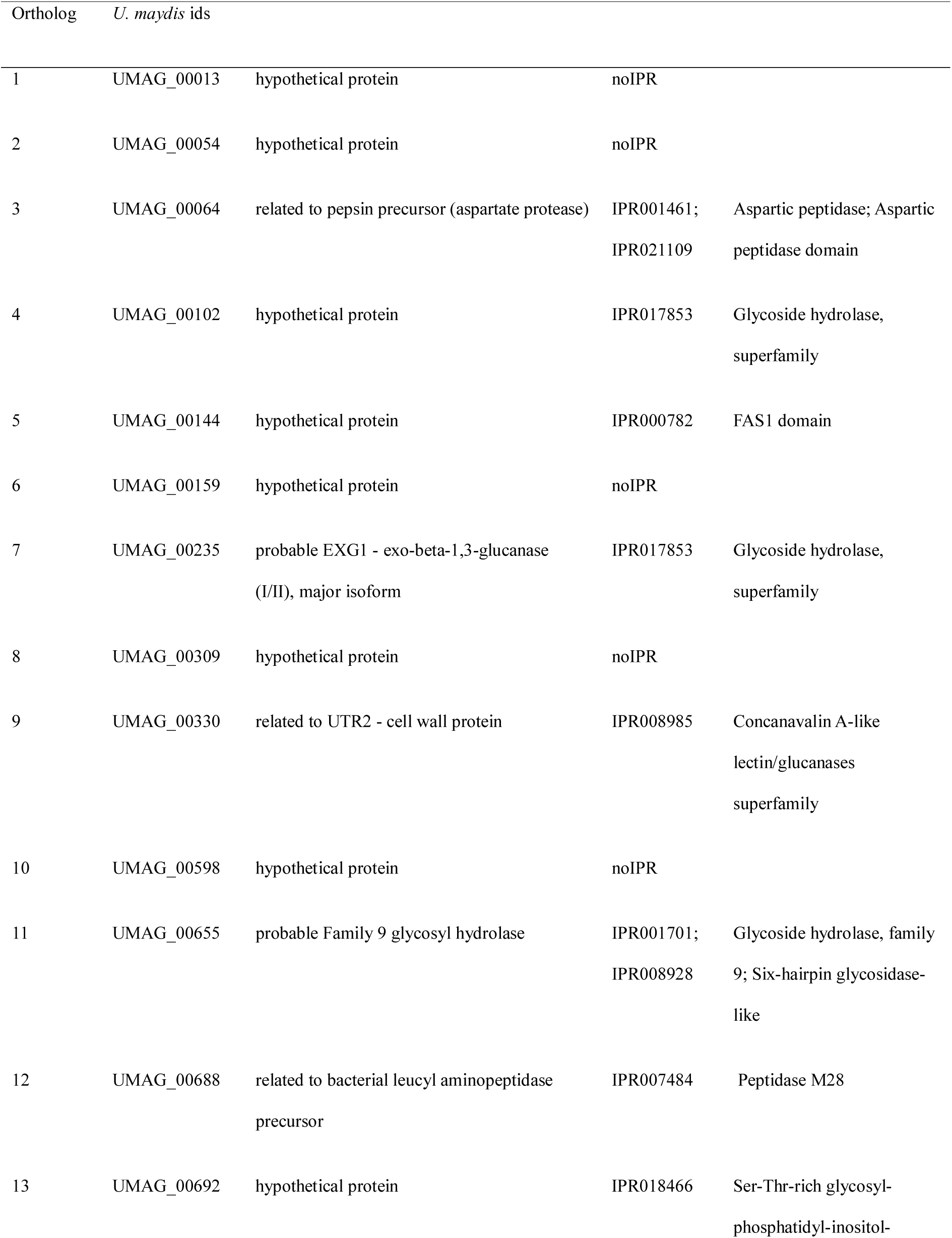
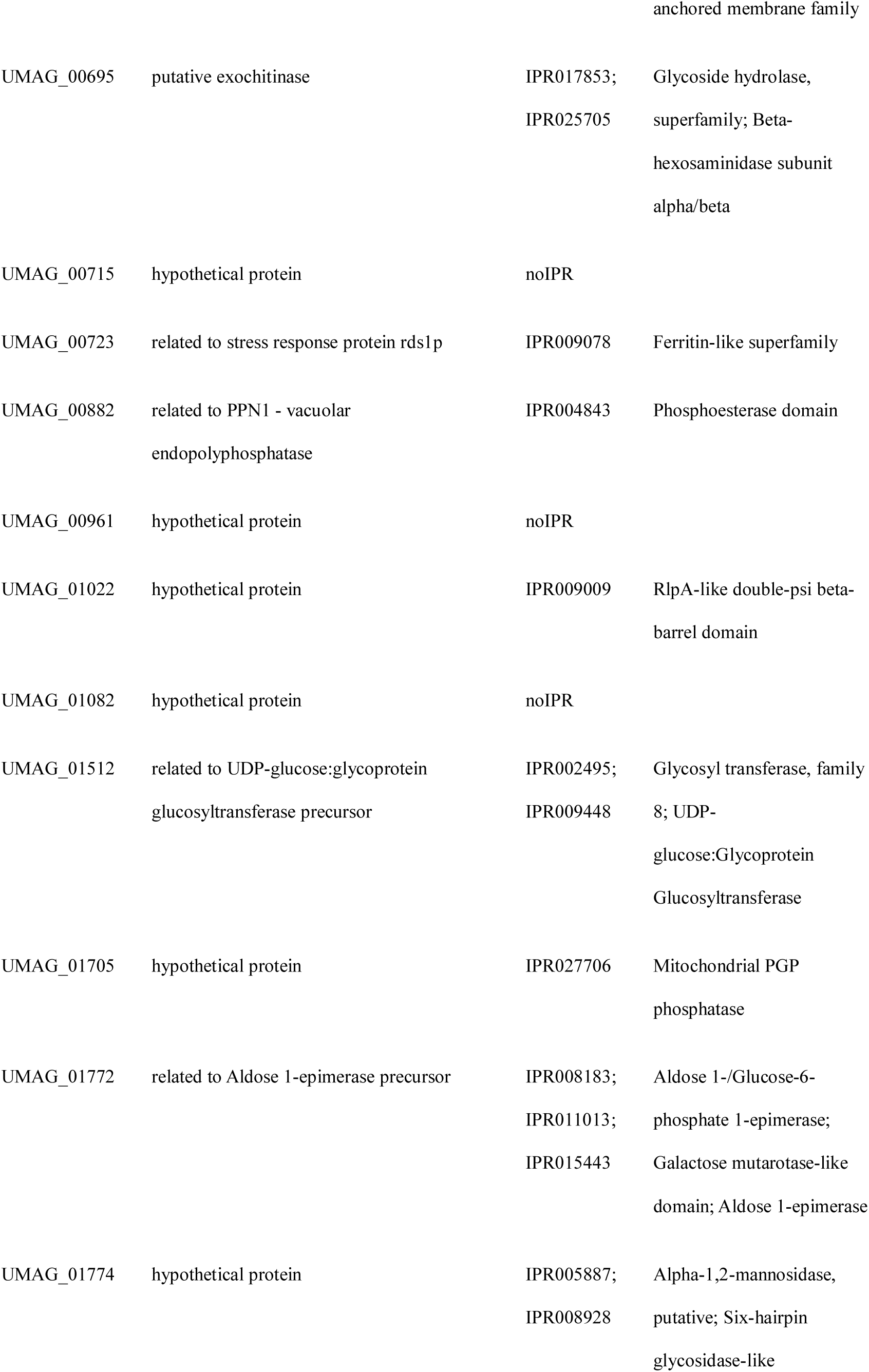
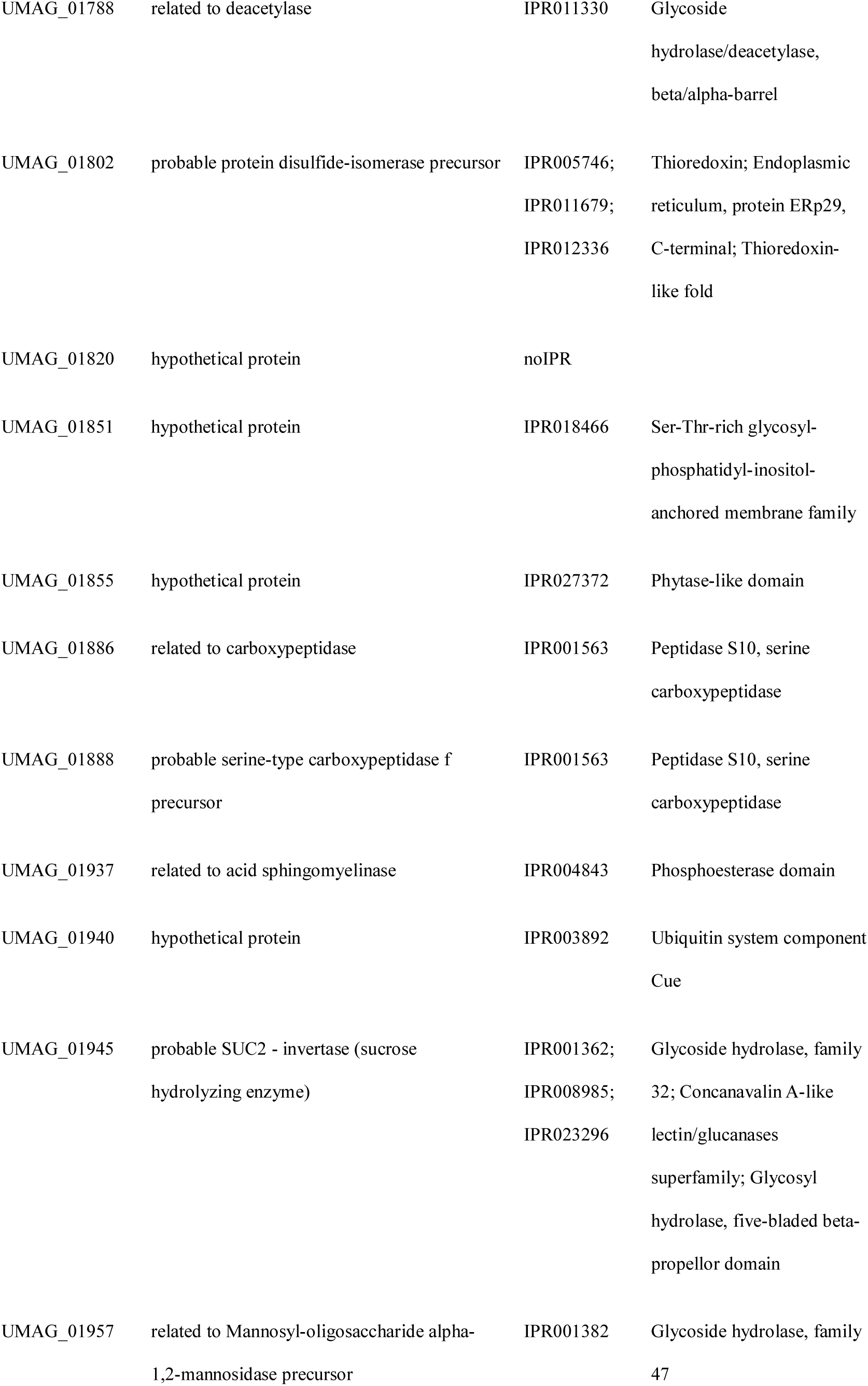
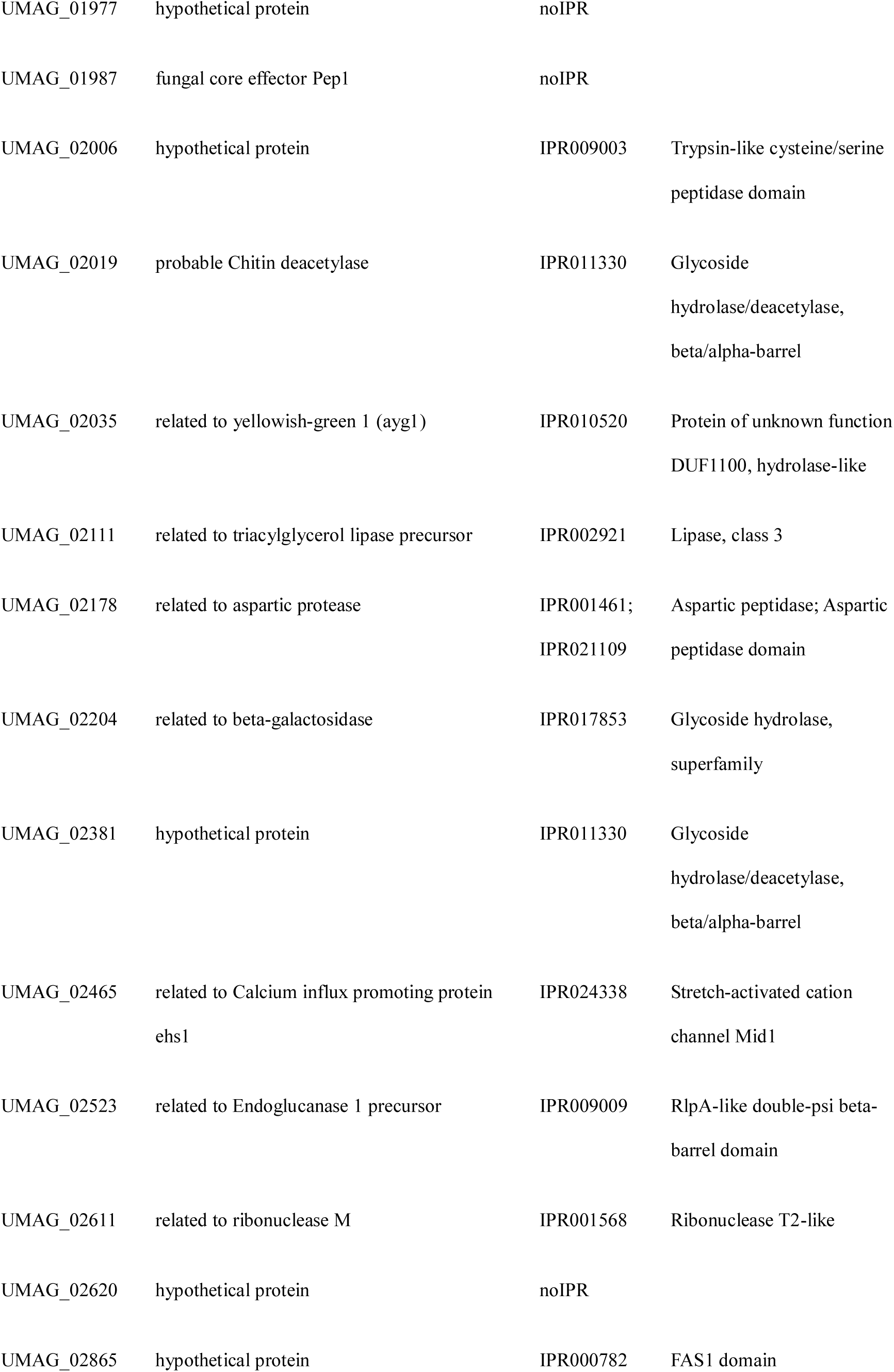
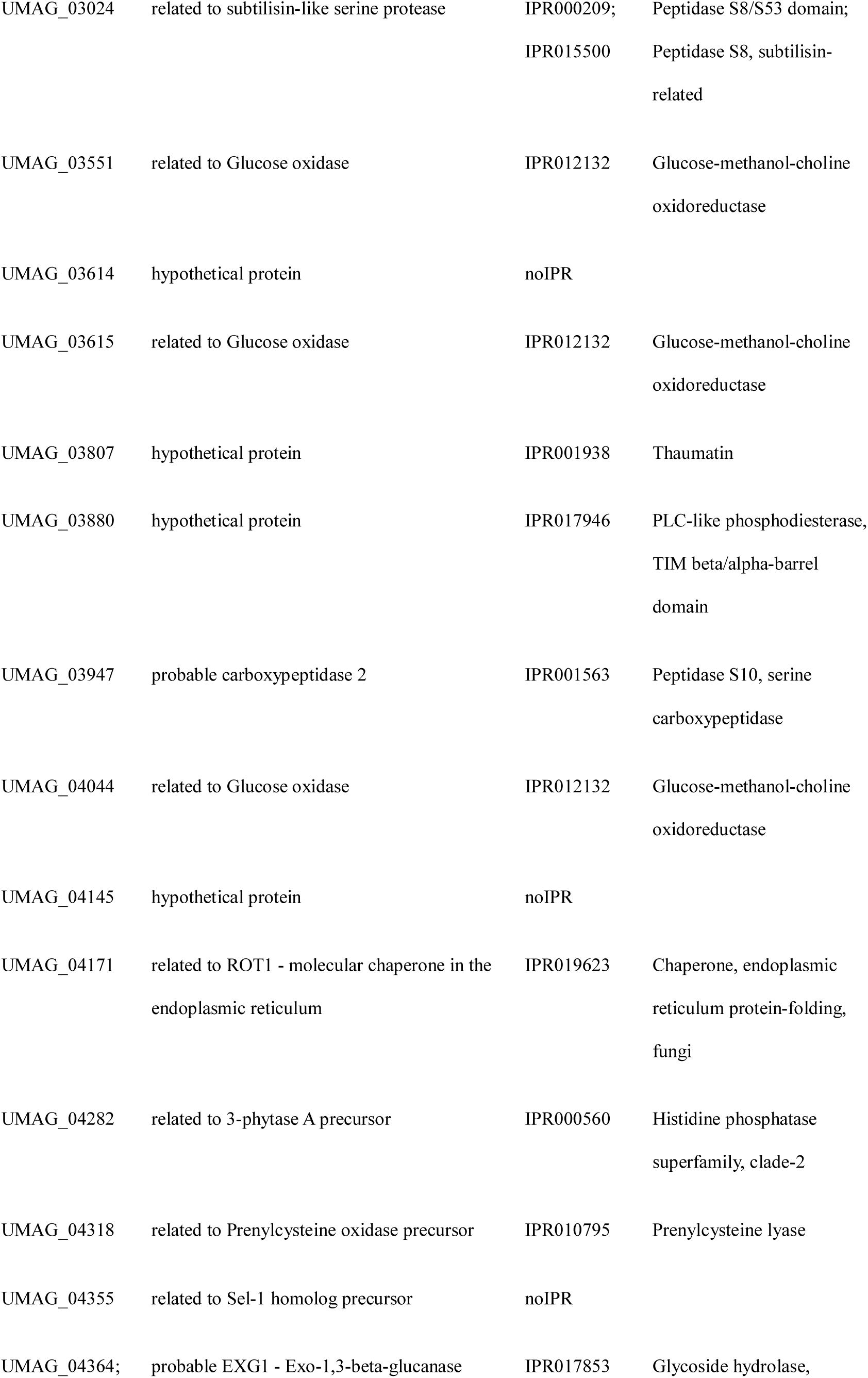
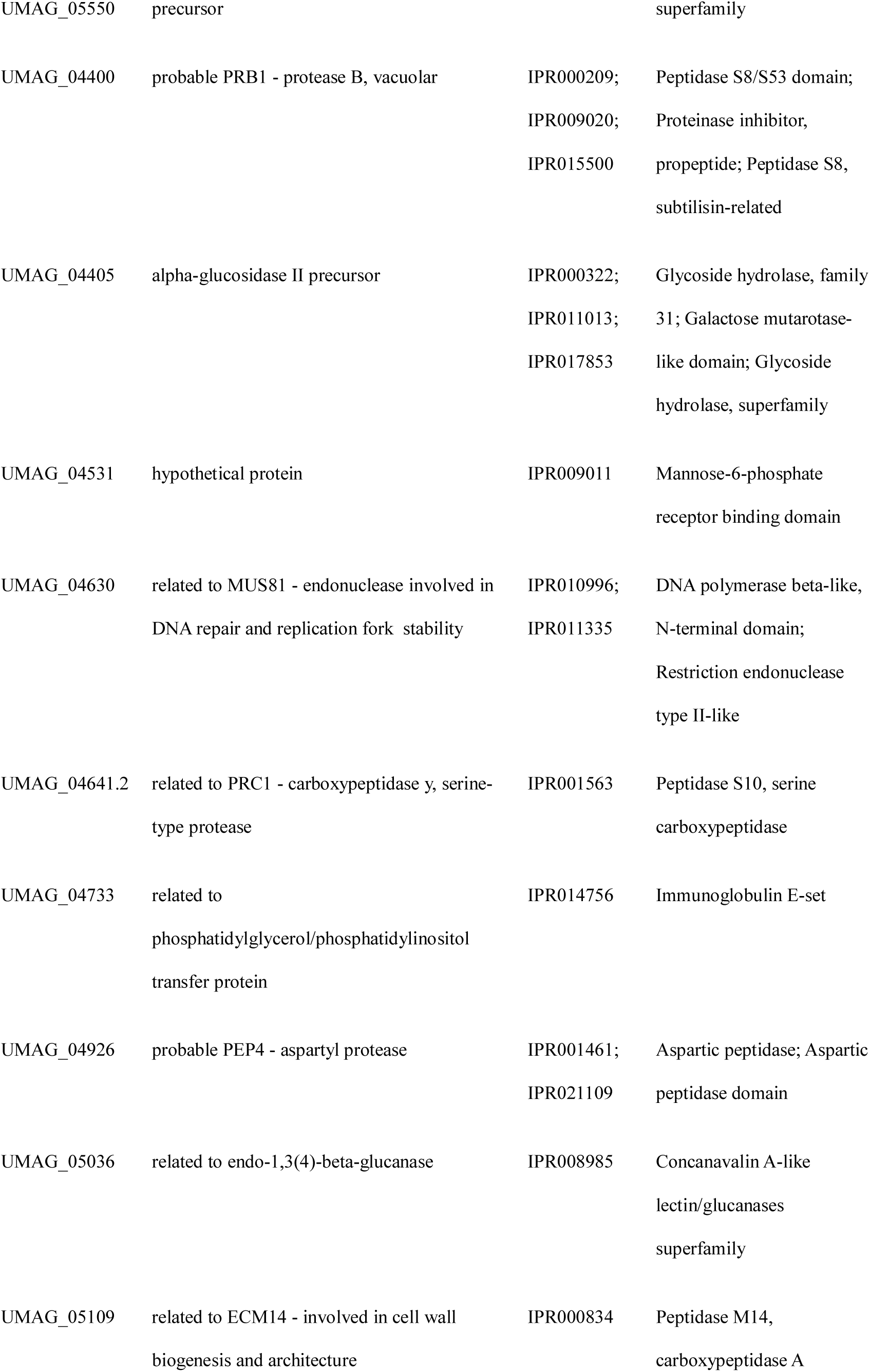
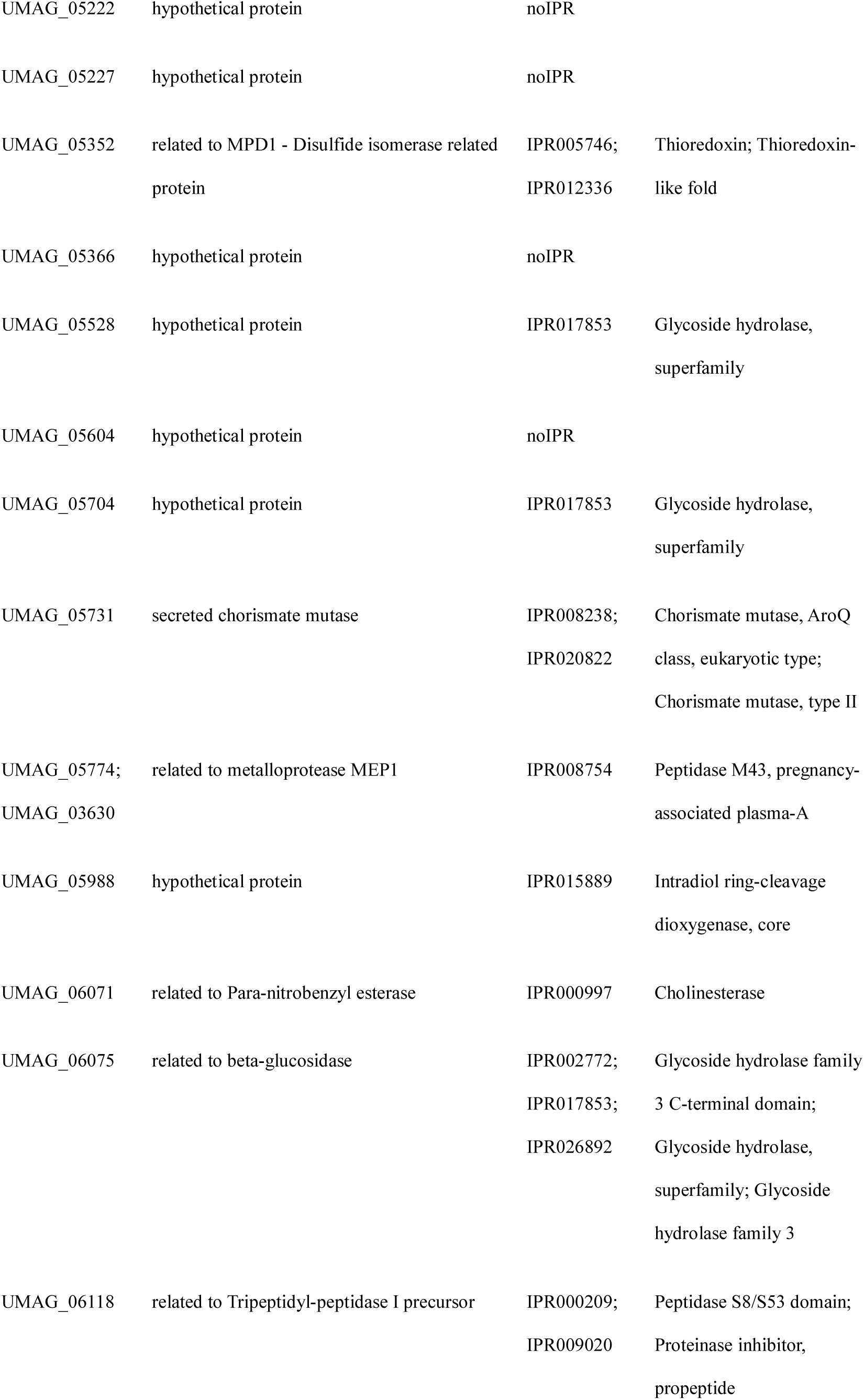
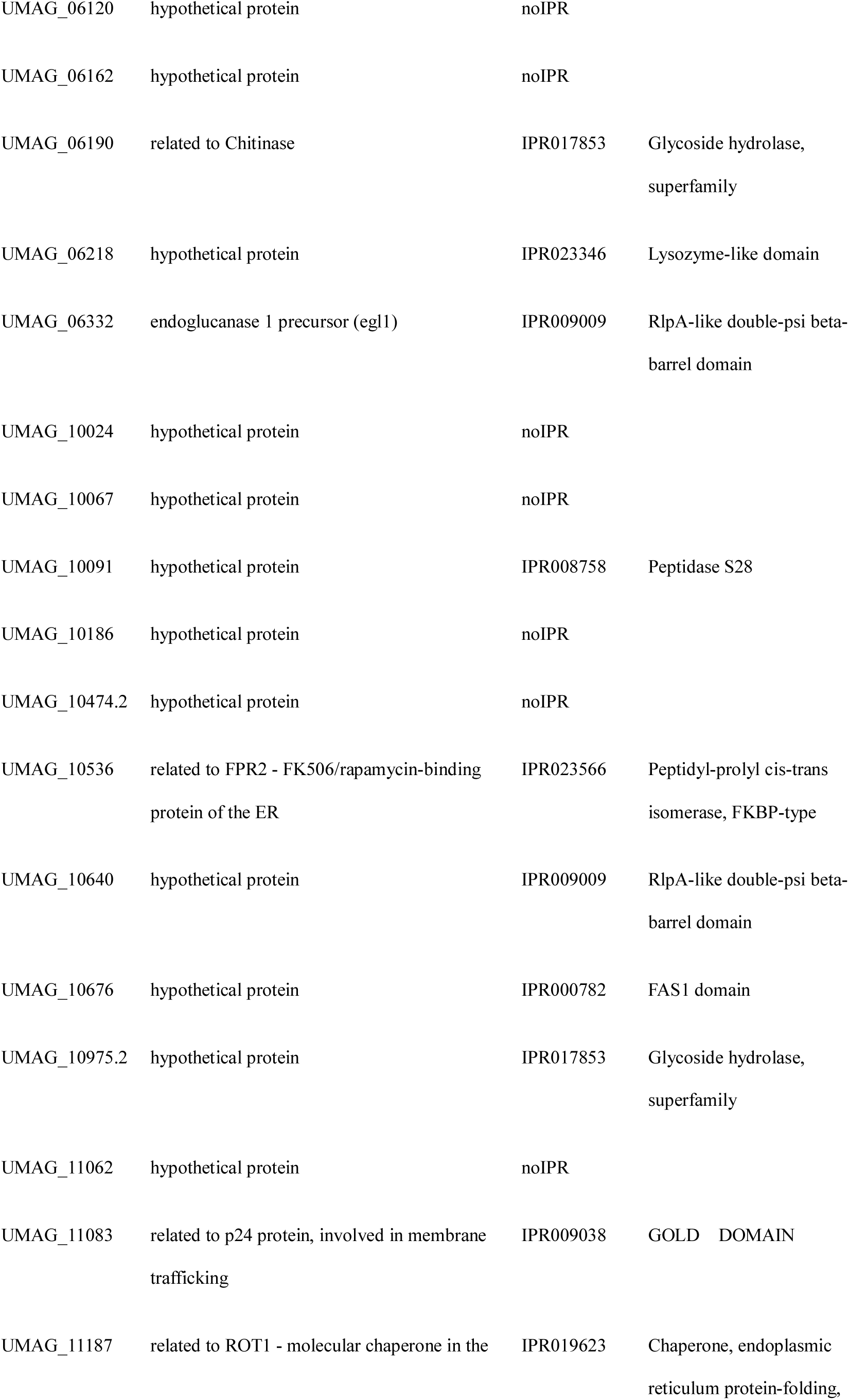
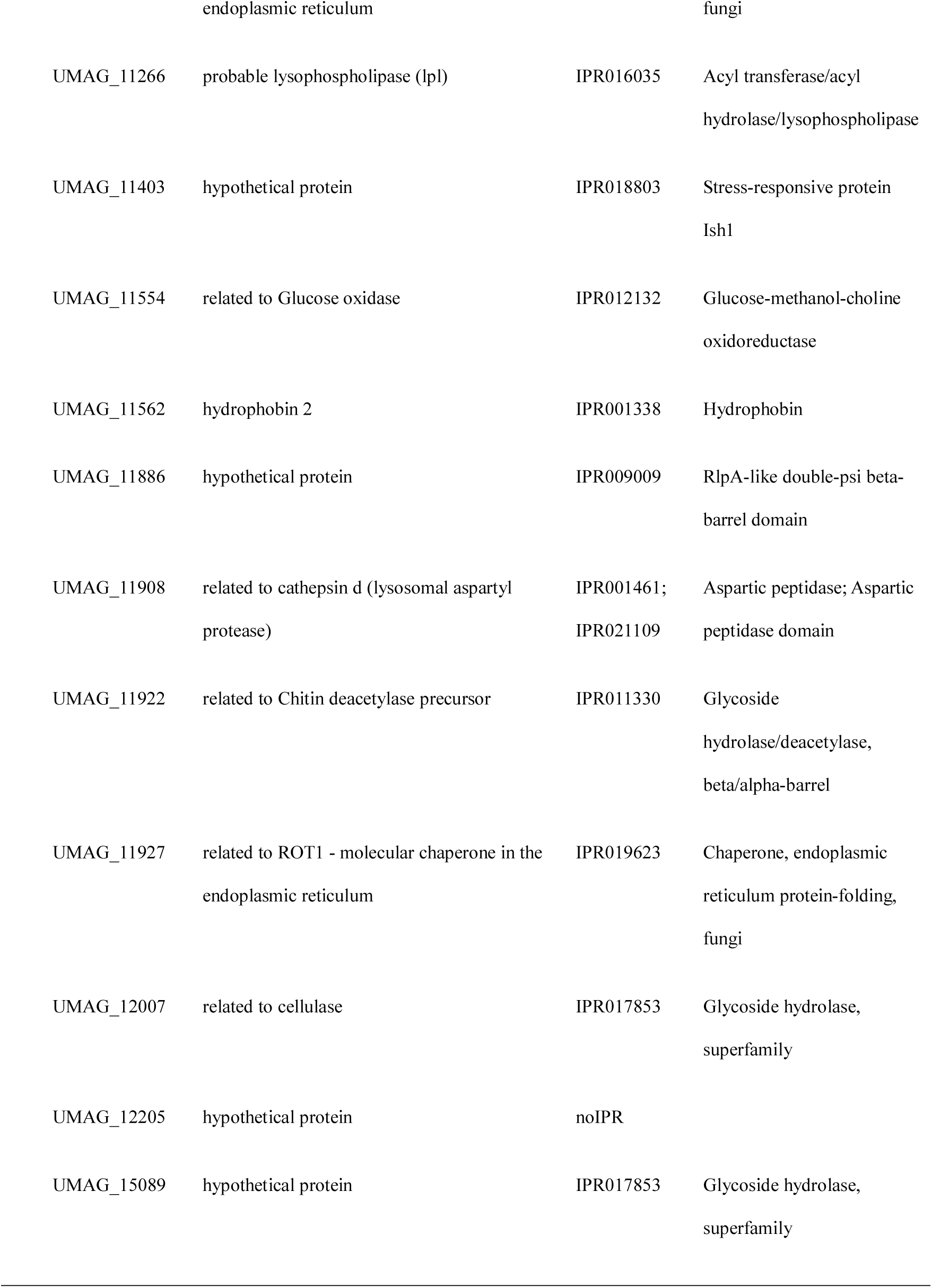
Annotation of putative secreted effector proteins conserved in all eight genomes.

