## Supplementary figures and images for "*Pseudozyma* saprotrophic yeasts have retained a large effector arsenal, including functional Pep1 orthologs"

### Supplementary file 1

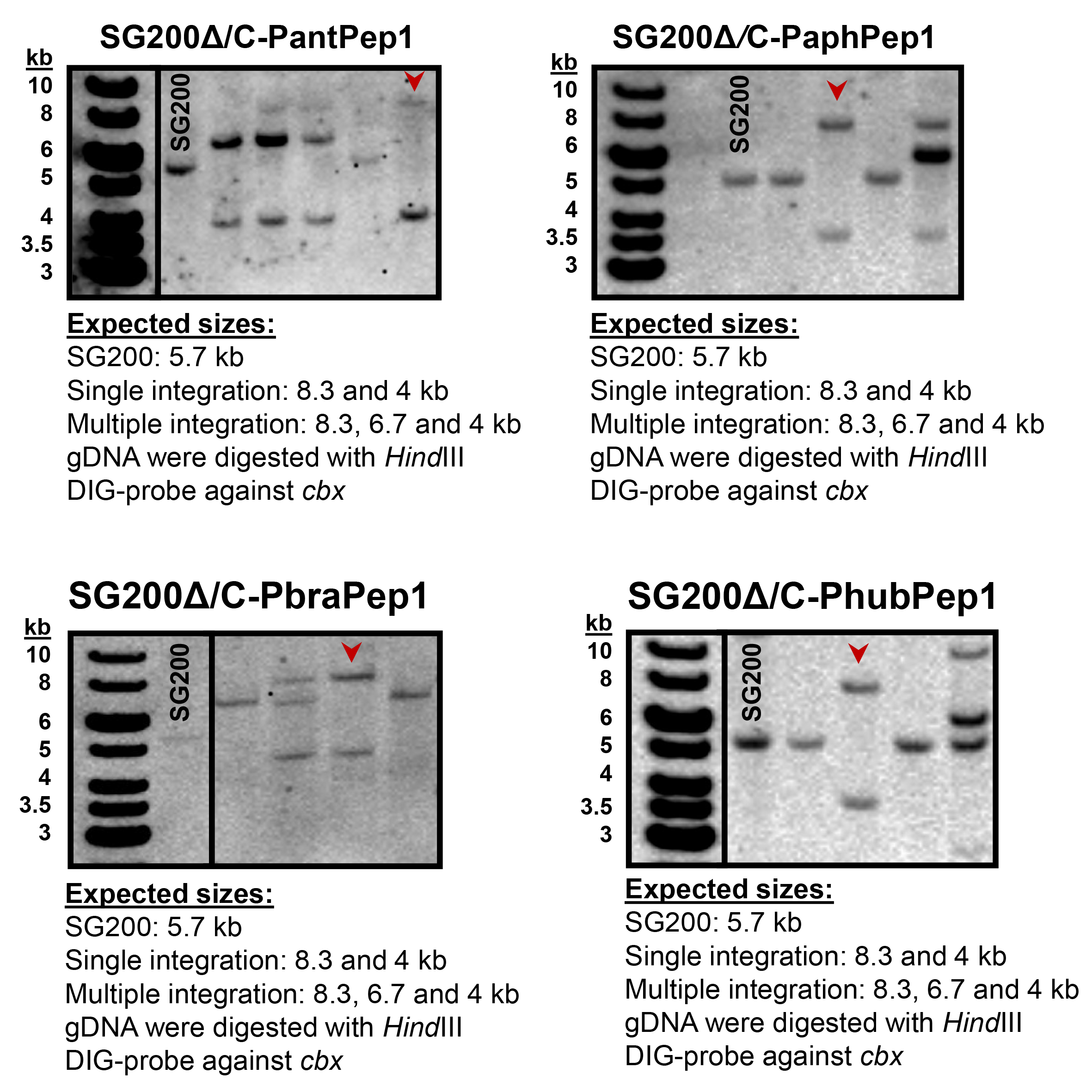
