## Supplementary material for "*Pseudozyma* saprotrophic yeasts have retained a large effector arsenal, including functional Pep1 orthologs"

Article history

This manuscript has first been submitted for publication in summer 2017. While the main conclusions have stayed the same – Pseudozyma species known only from the yeast stage have a hidden pathogenic stage – claims have been softened due to the criticism of reviewers. None of the reviewers had questioned the validity of the results, but equally none believed in the conclusions deduced form them, which is the reason why the current version of the manuscript contains more explanation and discussion on this topic.
